# Non-canonical HIPPO-MST1/2 promotes hyper-proliferation of pulmonary vascular cells through CDC20

**DOI:** 10.1101/2025.04.03.646845

**Authors:** Tapan Dey, Iryna Zhyvylo, Lifeng Jiang, Samuel O. Olapoju, Andressa Pena, Theodore Avolio, Derek Lin, Dmitry Goncharov, John R. Greenland, Paul J. Wolters, Horace DeLisser, Soni Savai Pullamsetti, Tatiana V. Kudryashova, Elena A. Goncharova

## Abstract

HIPPO components mammalian Ste20-like protein kinases 1 and 2 (MST1/2) are well described growth suppressors. However, in pulmonary arterial hypertension (PAH), MST1/2 switch their roles and become pro-proliferative and pro-survival molecules, supporting hyper-proliferation of pulmonary artery (PA) smooth muscle cells (PASMCs) and adventitial fibroblasts (PAAFs), remodeling of small PAs, and pulmonary hypertension. Here, we report that MST1/2 promotes hyper-proliferation and apoptosis resistance of human PAH PASMCs and PAAFs by up-regulating cell division cycle protein 20 (CDC20), establishing novel link between HIPPO-MST1/2 and cell cycle regulation in PAH.

**Authors Contributions:** conception and design of the work (EAG, SSP, TVK); acquisition, analysis, and interpretation of data (TD, IZ, LJ, SOO, AP, TA, DL, DG, JRG, PJW, HD, TK); drafting and editing the manuscript (EAG, SSP, TVK, JRG, PJW).

## To the Editor

Hyper-proliferation and resistance to apoptosis of pulmonary vascular cells in small muscular pulmonary arteries (PA) are important components of pulmonary vascular remodeling, a major irreversible feature of pulmonary arterial hypertension (PAH) (1). Components of HIPPO signaling mammalian Ste20-like protein kinases 1 and 2 (MST1/2) (also known as serine/threonine-protein kinases 4/3) act as growth suppressors in somatic mammalian cells (2). However, in PAH pulmonary vasculature, MST1/2 promote hyper-proliferation and apoptosis resistance of pulmonary artery (PA) smooth muscle cells (PASMCs) and adventitial fibroblasts (PAAFs) through up-regulating Budding Uninhibited By Benzimidazoles 3 Homolog (BUB3) and down-regulating Forkhead Box 3 (FOXO3), respectively (3). However, it is not clear whether PASMCs and PAAFs in PAH share any functional downstream effectors of MST1/2, which could be viewed as potential molecular targets for therapeutic intervention.

The cell division cycle protein 20 (CDC20) is a pro-proliferative protein acting as an oncogene in several human cancers (4). STRING functional protein association network database predicted that CDC20 connects BUB3, STK4 (MST1), STK3 (MST2), and FOXO3, (Figure 1A), suggesting its potential role as a downstream effector of MST1/2 that is shared between PASMCs and PAAFs. Analysis of publicly available databases GSE117261, GSE144274, and GSE144932 (5-7) demonstrated that *CDC20* expression is increased in the lungs and isolated PASMCs and PAAFs from patients with PAH compared to non-diseased controls (Figure 1B-D) and identified *CDC20* as a most significantly up-regulated gene in human PAH PASMCs (5).

**Figure 1.**
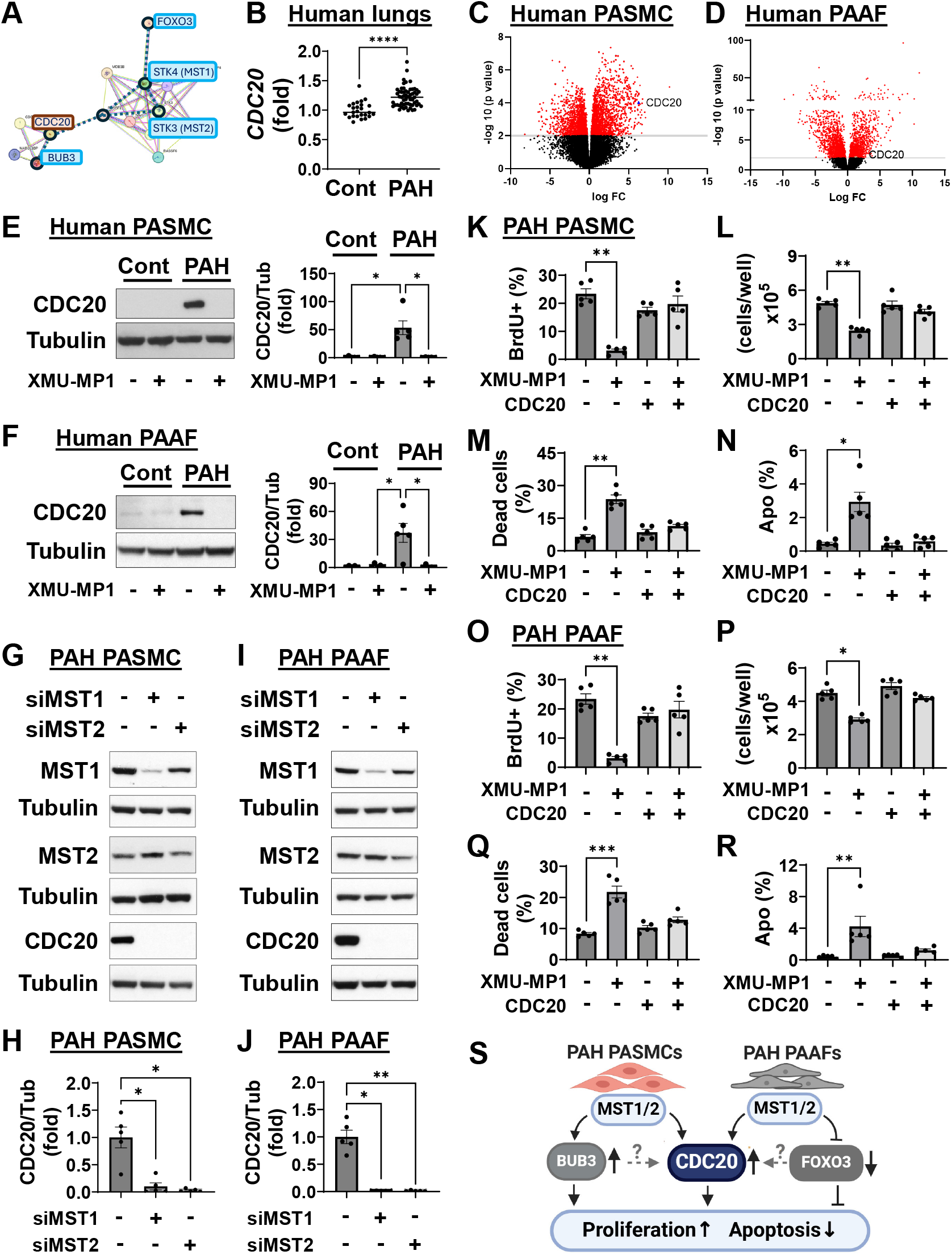
MST1/2 promotes hyper-proliferation and survival of human PAH PASMCs and PAAFs via CDC20. **A**: Predicted interactions among STK4 (MST1), STK3 (MST2), BUB3, FOXO3, and CDC20 by the STRING functional protein association network database. **B**: CDC20 gene expression in human lung tissue from non-diseased and PAH subjects. ****p<0.0001 by Mann Whitney U. Data are derived from publicly available dataset GSE117261. **C, D:** Volcano plots of differentially expressed genes (DEGs) in PASMCs (**C**) and PAAFs (**D**) from PAH vs control subjects. Data are derived from the publicly available datasets GSE144274 and GSE144932. **E, F**: Human control and PAH PASMCs (**E**) and PAAFs (**F**) were treated with diluent (−) or 5μM MST1/2 inhibitor XMU-MP1 for 48 hours followed by immunoblot analysis to detect indicated proteins. Data are means±SE; n=5 subjects/group. *p<0.05, **p<0.01 by Kruskal Wallis (Dunn post-hoc correction). **G-J**: Human PAH PASMCs (**G, H**) or PAAFs (**I, J**) were transfected with siRNA MST1, siRNA MST2, or control scrambled siRNA (−) for 48 hours followed by immunoblot analysis to detect indicated proteins. Representative images and data presented as means±SE from n=5 subjects/group. *p<0.05, **p<0.01 by Kruskal Wallis (Dunn post-hoc correction). **K-R**: Human PAH PASMCs (**K-N**) and PAAFs (**O-R**) were transfected with pCMV3-CDC20-GFPSpark or control pCMV3-GFPSpark plasmid (−) and treated with 5μM XMU-MP1 or diluent. 48 hours later, BrdU incorporation (**K, O**), cell counts (**L, P**), cell viability (% of dead cells) (**M, Q**) were performed; apoptosis was examined using *In situ* cell death detection assay (**N, R**). Data are means±SE from n=5 subjects/group. *p<0.05, **p<0.01, ***p<0.001 by Kruskal Wallis (Dunn post-hoc correction). **S**: MST1/2 promotes hyper-proliferation and survival of PASMCs and PAAFs via up-regulating CDC20. Further studies are needed to establish whether MST1/2 regulate CDC20 via BUB3 and FOXO3.

To test whether CDC20 acts downstream of MST1/2, we used pharmacological and molecular approaches. First, we treated early-passage human PAH and non-diseased (control) PASMCs and PAAFs with the selective MST1/2 inhibitor XMU-MP1. PASMCs and PAAFs from PAH patients had higher protein levels of CDC20 than respective control cells, which was significantly reduced by XMU-MP1 (Figure 1E, F). Supporting inhibitor-based findings, transfection of human PAH PASMCs and PAAFs with siRNA MST1 or siRNA MST2 dramatically reduced CDC20 protein content compared to cells transfected with control siRNA (Figure 1G-J). Taken together, these data show that CDC20 is overexpressed and acts as a downstream positive effector of MST1/2 in PASMCs and PAAFs from patients with PAH.

To explore potential role of CDC20 in MST1/2-induced hyper-proliferation and survival of PAH pulmonary vascular cells, we used a “rescue” approach. We transfected human PAH PASMCs and PAAFs with mammalian vector expressing human CDC20 (pCMV3-CDC20-GFPSpark) or empty pCMV3-GFPSpark plasmid (control) in the presence or absence of MST1/2 inhibitor XMU-MP1 and tested cell proliferation and survival using BrdU incorporation, cell count and viability assays, and *In Situ* Cell Death Detection Kit (3, 8). In line with published findings (3), XMU-MP1 significantly reduced proliferation and induced apoptosis in PAH PASMCs and PAAFs transfected with control plasmid (Figure 1K-R). In contrast, transfection with pCMV3-CDC20-GFPSpark protected cells from these XMU-MP1 effects (Figure 1K-R), suggesting that MST1/2 promote proliferation and survival of human PAH PASMCs and PAAFs via CDC20.

In conclusion, our data identified CDC20 as a novel downstream effector of non-canonical HIPPO-MST1/2 signaling, which is required for MST1/2-dependent hyper-proliferation and apoptosis resistance of PASMCs and PAAFs from human PAH lungs (Figure 1S), providing novel link between HIPPO cassette and cell cycle regulation. Further studies are needed to determine the relationship between CDC20 and previously identified PAH-specific MST1/2 downstream effectors BUB3 and FOXO3. It is also important to note that dysregulation of cell cycle components cyclin-dependent kinases (CDKs) has been recently reported as an important driver of pulmonary vascular hyperproliferation, remodeling, and PAH (9). CDKs and CDC20 reciprocally regulate each other to drive cell cycle in physiological conditions (10). Our findings raise the possibility that MST1/2-induced CDC20 up-regulation could disturb physiological CDC20-CDKs coordination, further supporting unstimulated pulmonary vascular hyper-proliferation and remodeling in PAH. Our findings strongly suggest that further studies are needed to determine the mechanisms of regulation and function of CDC20 in healthy and diseased pulmonary vasculature and to evaluate a potential attractiveness of targeting CDC20 as a remodeling-focused strategy to treat PAH.

## Supporting information

Dey et al Online Supplement

## Notes

### Competing Interest Statement

The authors have declared no competing interest.

## References

1. Johnson S, Sommer N, Cox-Flaherty K, Weissmann N, Ventetuolo CE, Maron BA. Pulmonary Hypertension: A Contemporary Review. Am J Respir Crit Care Med 2023; 208:528–548.

2. Dey A, Varelas X, Guan KL. Targeting the Hippo pathway in cancer, fibrosis, wound healing and regenerative medicine. Nat Rev Drug Discov 2020; 19:480–494.

3. Kudryashova TV, Dabral S, Nayakanti S, Ray A, Goncharov DA, Avolio T, Shen Y, Rode A, Pena A, Jiang L, Lin D, Baust J, Bachman TN, Graumann J, Ruppert C, Guenther A, Schmoranzer M, Grobs Y, Eve Lemay S, Tremblay E, Breuils-Bonnet S, Boucherat O, Mora AL, DeLisser H, Zhao J, Zhao Y, Bonnet S, Seeger W, Pullamsetti SS, Goncharova EA. Noncanonical HIPPO/MST Signaling via BUB3 and FOXO Drives Pulmonary Vascular Cell Growth and Survival. Circ Res 2022; 130:760–778.

4. Bruno S, Ghelli Luserna di Rorà A, Napolitano R, Soverini S, Martinelli G, Simonetti G. CDC20 in and out of mitosis: a prognostic factor and therapeutic target in hematological malignancies. J Exp Clin Cancer Res 2022; 41:159.

5. Gorr MW, Sriram K, Muthusamy A, Insel PA. Transcriptomic analysis of pulmonary artery smooth muscle cells identifies new potential therapeutic targets for idiopathic pulmonary arterial hypertension. Br J Pharmacol 2020; 177:3505–3518.

6. Stearman RS, Bui QM, Speyer G, Handen A, Cornelius AR, Graham BB, Kim S, Mickler EA, Tuder RM, Chan SY, Geraci MW. Systems Analysis of the Human Pulmonary Arterial Hypertension Lung Transcriptome. Am J Respir Cell Mol Biol 2019; 60:637–649.

7. Chelladurai P, Kuenne C, Bourgeois A, Günther S, Valasarajan C, Cherian AV, Rottier RJ, Romanet C, Weigert A, Boucherat O, Eichstaedt CA, Ruppert C, Guenther A, Braun T, Looso M, Savai R, Seeger W, Bauer UM, Bonnet S, Pullamsetti SS. Epigenetic reactivation of transcriptional programs orchestrating fetal lung development in human pulmonary hypertension. Sci Transl Med 2022; 14:eabe5407.

8. Goncharov DA, Kudryashova TV, Ziai H, Ihida-Stansbury K, DeLisser H, Krymskaya VP, Tuder RM, Kawut SM, Goncharova EA. Mammalian target of rapamycin complex 2 (mTORC2) coordinates pulmonary artery smooth muscle cell metabolism, proliferation, and survival in pulmonary arterial hypertension. Circulation 2014; 129:864–874.

9. Weiss A, Neubauer MC, Yerabolu D, Kojonazarov B, Schlueter BC, Neubert L, Jonigk D, Baal N, Ruppert C, Dorfmuller P, Pullamsetti SS, Weissmann N, Ghofrani HA, Grimminger F, Seeger W, Schermuly RT. Targeting cyclin-dependent kinases for the treatment of pulmonary arterial hypertension. Nat Commun 2019; 10:2204.

10. Hein JB, Nilsson J. Interphase APC/C-Cdc20 inhibition by cyclin A2-Cdk2 ensures efficient mitotic entry. Nat Commun 2016; 7:10975.

