## Supplementary material for "Non-canonical HIPPO-MST1/2 promotes hyper-proliferation of pulmonary vascular cells through CDC20": Dey et al Online Supplement

**Supplemental Materials and Methods**

**Human cell cultures**

Human lung tissues for cell isolation and early-passage (3–8 passage) human pulmonary arterial (PA) smooth muscle cells (PASMCs) and pulmonary artery adventitial fibroblasts (PAAFs) isolated from small (<1.5 mm outer diameter) PAs of non-diseased subjects (postmortem) and subjects with pulmonary arterial hypertension (PAH) were provided by University of Pittsburgh VMI Cell Processing Core, UC Davis Lung Center Pulmonary Vascular Disease Program human specimens biobank, UC San Francisco transplant program, and the Pulmonary Hypertension Breakthrough Initiative (PHBI) under approved protocols in accordance with Institutional Review Boards and Committee for Oversight of Research and Clinical Training Involving Decedents. Please see Supplemental Table 1 for human subjects’ characteristics. Cells isolation, characterization and maintenance were performed under protocols approved by PHBI as described previously (1-4). Cells were grown in complete Smooth Muscle Cell Growth Medium 2 (PASMCs) (Cat#C-22162, PromoCell, Heidelberg, Germany) or complete Fibroblasts Growth Medium 2 (Cat#C-23120, PromoCell) and Antibiotic-Antimycotic (Cat#15240062, Thermo Fisher Scientific, Waltham, MA). Before experiments, cells were maintained in the Smooth Muscle Cell basal medium 2 (Cat#C-22262, PromoCell) (PASMCs) or Fibroblast basal medium 2 (Cat#C-23220, PromoCell) (PAAFs) supplemented with 5% fetal bovine serum (FBS) and serum-deprived for 48 hours in basal media supplemented with 0.1% bovine serum albumin (BSA).

**Immunoblot analysis** was performed as previously described (1-4). XMU-MP1 was purchased from Bio Techne-Sales Corporation (Minneapolis, MA) (Cat#6482/10). Primary antibodies for MST1 (Cat#3682) (1:1000 dilution), MST2 (Cat#3952) (1:1000 dilution), CDC20 (Cat#4823) (1:1000 dilution), tubulin (Cat#2148) (1:2000 dilution), and HRP-linked anti-rabbit IgG (Cat#7074) (1:1000 – 1:2500 dilution) were purchased from Cell Signaling (Danvers, MA). Each experiment was repeated five times, cells from different human subjects were used in each experiment.

**Cell Transfection** was performed as described previously (1, 4). Silencer^TM^ Select siRNA STK4 (MST1) (Cat#4390825; siRNA Assay ID# S13570) and Silencer^TM^ Select siRNA STK3 (MST2) (Cat#4390825; siRNA Assay ID #S13567) were purchased from Life Technologies Corporation (Carlsbad, CA). ON TARGETplus non-targeting control siRNA (Cat#D-001810-01-20) was purchased from Horizon Discovery Bioscience Limited (Cambridge, UK). Mammalian vector pCMV3-C-CDC20-GFPSpark (Cat#HG16751-ACG) and control pCMV3-C-GFPSpark plasmids (Cat#CV026**)** were purchased from Genescript USA Inc (Piscataway, NJ). Transfections of siRNAs were performed using DharmaFECT 1 Transfection Reagent (Cat#T-2001-03, Horizon Discovery Bioscience Limited); transfections of plasmids were performed using Effectene Transfection Reagent (Cat#301427, Qiagen Inc, Germantown, MD) according to the manufacturers’ protocols.

**Proliferation and Apoptosis Assays**

BrdU incorporation, cell counts, and cell viability assays were performed as described previously (1, 3, 4). BrdU monoclonal antibody (Cat# B35128) were purchased from Thermo Fisher Scientific. Cell counts and viability were measured using Countess 3 automated Cell Counter (Thermo Fisher Scientific). Apoptosis was evaluated using *In Situ* Cell Death Detection Kit, Fluorescein (Cat#11684795910, Millipore Sigma, Sant Luise, MO) following manufacturer’s protocol as previously described (2-4). For BrdU and apoptosis analyses, DAPI (Cat#D21490, Thermo Fisher Scientific) staining was used to detect nuclei. Images were taken using Keyence BZX810 microscope (Keyence, Osaka, Japan). For cell count and cell viability assays, three technical repetitions per condition per experiment were performed. For BrdU incorporation and apoptosis experiments, minimum of 200 cells/condition/experiment were analyzed, the percentage of BrdU-positive or TUNEL-positive cells per total number of cells was calculated. For each cell type and each assay, five independent experiments were conducted, each experiment was performed using cells from different human subjects.

**Data Analysis**

In order to reduce bias, blinded data analysis was performed when possible. Immunoblot images were analyzed using ImageJ (National Institutes of Health, Bethesda, MD, United States). Statistical analysis was performed using GraphPad Prizm 9.02 (GraphPad Software, San Diego, CA, United States). Statistical comparisons between the two groups were performed by Mann-Whitney U test. Statistical comparisons among three and more groups were performed by Kruskal Wallis test with Dunn multiple comparison test. Statistical significance was defined as p≤ 0.05.

**Supplemental Tables**

**Table 1.** **Human subjects’ characteristics**

| Diagnosis | Gender | Age (Years) |
| --- | --- | --- |
| Non-diseased | F | 44 |
| Non-diseased | F | 38 |
| Non-diseased | M | 19 |
| Non-diseased | M | 47 |
| Non-diseased | M | 70 |
| Non-diseased | M | 36 |
| Non-diseased | F | 38 |
| Non-diseased | F | 64 |
| Non-diseased | F | 48 |
| Non-diseased | F | 43 |
| IPAH | F | 40 |
| IPAH-Sc | F | 53 |
| IPAH | F | 39 |
| IPAH | F | 62 |
| IPAH | F | 43 |
| IPAH | F | 50 |
| IPAH | M | 21 |
| IPAH | F | 36 |

IPAH – idiopathic pulmonary arterial hypertension; Sc- scleroderma; F – female; M - male

**Supplemenral References**

1. Goncharov DA, Kudryashova TV, Ziai H, Ihida-Stansbury K, DeLisser H, Krymskaya VP, Tuder RM, Kawut SM, Goncharova EA. Mammalian target of rapamycin complex 2 (mTORC2) coordinates pulmonary artery smooth muscle cell metabolism, proliferation, and survival in pulmonary arterial hypertension. *Circulation* 2014; 129: 864-874.

2. Jiang L, Goncharov DA, Shen Y, Lin D, Chang B, Pena A, DeLisser H, Goncharova EA, Kudryashova TV. Akt-Dependent Glycolysis-Driven Lipogenesis Supports Proliferation and Survival of Human Pulmonary Arterial Smooth Muscle Cells in Pulmonary Hypertension. *Front Med (Lausanne)* 2022; 9: 886868.

3. Shen Y, Goncharov DA, Pena A, Baust J, Chavez Barragan A, Ray A, Rode A, Bachman TN, Chang B, Jiang L, Dieffenbach P, Fredenburgh LE, Rojas M, DeLisser H, Mora AL, Kudryashova TV, Goncharova EA. Cross-talk between TSC2 and the extracellular matrix controls pulmonary vascular proliferation and pulmonary hypertension. *Sci Signal* 2022; 15: eabn2743.

4. Kudryashova TV, Dabral S, Nayakanti S, Ray A, Goncharov DA, Avolio T, Shen Y, Rode A, Pena A, Jiang L, Lin D, Baust J, Bachman TN, Graumann J, Ruppert C, Guenther A, Schmoranzer M, Grobs Y, Eve Lemay S, Tremblay E, Breuils-Bonnet S, Boucherat O, Mora AL, DeLisser H, Zhao J, Zhao Y, Bonnet S, Seeger W, Pullamsetti SS, Goncharova EA. Noncanonical HIPPO/MST Signaling via BUB3 and FOXO Drives Pulmonary Vascular Cell Growth and Survival. *Circ Res* 2022; 130: 760-778.
